# Iso-Seq allows genome-independent transcriptome profiling of grape berry development

**DOI:** 10.1101/269530

**Authors:** Andrea Minio, Mélanie Massonnet, Rosa Figueroa-Balderas, Amanda M. Vondras, Barbara Blanco-Ulate, Dario Cantu

**Affiliations:** Department of Viticulture and Enology, University of California Davis, Davis, CA; Department of Plant Sciences, University of California Davis, Davis, CA

**Keywords:** transcriptome reference, RNA-seq, *Vitis vinifera*, berry ripening, alternative splicing

## Abstract

Transcriptomics has been widely applied to study grape berry development. With few exceptions, transcriptomic studies in grape are performed using the available genome sequence, PN40024, as reference. However, differences in gene content among grape accessions, which contribute to phenotypic differences among cultivars, suggest that a single reference genome does not represent the species’ entire gene space. Though whole genome assembly and annotation can reveal the relatively unique or “private” gene space of any particular cultivar, transcriptome reconstruction is a more rapid, less costly, and less computationally intensive strategy to accomplish the same goal. In this study, we used single molecule-real time sequencing (Iso-Seq) to sequence full-length cDNA and reconstruct the transcriptome of Cabernet Sauvignon berries during berry ripening. In addition, Illumina short reads from ripening berries were used to error-correct low-expression isoforms and to profile isoform expression. By comparing the annotated gene space of Cabernet Sauvignon to other grape cultivars, we demonstrate that the transcriptome reference built with Iso-Seq data represents most of the expressed genes in the grape berries and includes 1,501 cultivar-specific genes. Iso-Seq produced transcriptome profiles similar to those obtained after mapping on a complete genome reference. Together, these results justify the application of Iso-Seq to identify cultivar-specific genes and build a comprehensive reference for transcriptional profiling that circumvents the necessity of a genome reference with its associated costs and computational weight.

## INTRODUCTION

Grape berries undergo a series of complex physiological and biochemical changes during their development that determine their characteristics at harvest (Kuhn *et al.* 2014). Genome-wide expression studies using microarray and, more recently, RNA sequencing (RNA-seq) revealed that berry development involves the expression and modulation of approximately 23,000 genes (Massonnet *et al.* 2017a) and that the ripening transition is associated with a major transcriptome shift (Fasoli *et al.* 2012). Transcriptomic studies characterized the ripening program across grapevine cultivars (Venturini *et al.* 2013; Da Silva *et al.* 2013; Jiao *et al.* 2015; Massonnet *et al.* 2017a), identifying key ripening related genes (Palumbo *et al.* 2014; Massonnet *et al.* 2017a) and determining the impact of stress and viticultural practices on ripening (Deluc *et al.* 2009; Pastore *et al.* 2013; Xi *et al.* 2014; Amrine *et al.* 2015; Blanco-Ulate *et al.* 2015, 2017; Corso *et al.* 2015; Hopper *et al.* 2016; Savoi *et al.* 2016; Zenoni *et al.* 2017; Lecourieux *et al.* 2017; Massonnet *et al.* 2017b). This knowledge increases the possibility of exerting control over the ripening process, improving fruit composition under suboptimal or adverse conditions, and enhancing desirable traits in a crop with outstanding cultural and commercial significance (Savoi *et al.* 2016, 2017; Serrano *et al.* 2017; Zenoni *et al.* 2017).

These genome-wide expression analyses were possible because a highly contiguous assembly for the species was produced (Jaillon *et al.* 2007); this first effort used a grape line (PN40024) created by several rounds of backcrossing to reduce heterozygosity, facilitating genome assembly (Jaillon *et al.* 2007). Though poor by current standards, this pioneering, chromosome-resolved assembly served as the basis for numerous publications. However, the structural diversity of grape genomes makes using a single one-size-fits-all reference genome inappropriate (Golicz *et al.* 2016b, 2016a). There is substantial unshared gene content between cultivars, with 8 - 10% of the genes missing when two cultivars are compared (Da Silva *et al.* 2013). Although many of these genes are not essential for plant survival, they can account for 80% of the expression within their respective families and expand key gene families possibly associated with cultivar-specific traits (Da Silva *et al.* 2013).

Assembling genome references for all interesting cultivars is impractical, in part because its cost remains prohibitive and because of genomic features that impede the development of high-quality genome assemblies for any grape cultivar. Although the *V. vinifera* genome is relatively small (~487Mb) (Lodhi and Reisch 1995; Jaillon *et al.* 2007) and as repetitive as other plant genomes of similar size (41.4%) (Jaillon *et al.* 2007; Michael and Jackson 2013), it is highly heterozygous (~13%) (Jaillon *et al.* 2007; Velasco *et al.* 2007). Most domesticated grape cultivars are crosses between distantly related parents; this and clonal propagation cause the high heterozygosity observed in the species (Strefeler *et al.* 1992; Ohmi *et al.* 1993; Bowers and Meredith 1997; Sefc *et al.* 1998; Lopes *et al.* 1999; Di Gaspero *et al.* 2005; Tapia *et al.* 2007; Ibáñez *et al.* 2009; Cipriani *et al.* 2010; Myles *et al.* 2011; Lacombe *et al.* 2013; Chin *et al.* 2016; Minio *et al.* 2017; Zhou *et al.* 2017). Earlier attempts using short reads struggled to resolve complex, highly heterozygous genomes (Gnerre *et al.* 2011; Huang *et al.* 2012; Di Genova *et al.* 2014; Kajitani *et al.* 2014; Safonova *et al.* 2015). A limited ability to call consensus polymorphic regions yields highly fragmented assemblies where structural ambiguity occurs and alternative alleles at heterozygous sites are excluded altogether (Velasco *et al.* 2007). Single Molecule Real Time (SMRT) DNA sequencing (Pacific Biosciences, California, USA) has emerged as the leading technology for reconstructing highly contiguous, diploid assemblies of long, repetitive genomes that include phased information about heterozygous sites (Chin *et al.* 2013, 2016; Doi *et al.* 2014; Gordon *et al.* 2016; Pryszcz and Gabaldón 2016; Ricker *et al.* 2016; Seo *et al.* 2016; Vij *et al.* 2016; Huddleston *et al.* 2017). Recently, we used *Vitis vinifera* cv. Cabernet Sauvignon to test the ability of SMRT reads and the FALCON-Unzip assembly pipeline to resolve both alleles at heterozygous sites in the genome (Chin *et al.* 2016). The assembly produced was significantly more contiguous (contig N50 = 2.17 Mb) than the original PN40024 assembly (contig N50 = 102.7 kbp) and provided the first phased sequences of the diploid *V. vinifera* genome (Minio *et al.* 2017).

Despite recent advances in genome reconstruction methodologies, assembling a complex plant genome is still costly. Transcriptome reconstruction is the only alternative strategy to depict known and unknown gene content information (Venturini *et al.* 2013; Da Silva *et al.* 2013; Jiao *et al.* 2015). *De novo* assembly of RNA-seq reads is widely used for this purpose (Grabherr *et al.* 2011; Ashrafi *et al.* 2012; Venturini *et al.* 2013; Bellucci *et al.* 2014). SMRT technology was recently deployed to investigate expressed gene isoforms (Iso-Seq) in a variety of organisms, including a handful of plant species (Liu *et al.* 2017; Zulkapli *et al.* 2017; Filichkin *et al.* 2018).

Long reads delivered by this methodology report full-length transcripts sequenced from their 5’-ends to polyadenylated tails (Dong *et al.* 2015; Weirather *et al.* 2015; Gao *et al.* 2016; Tombácz *et al.* 2016; Kuo *et al.* 2017; Price and Gibas 2017; Workman *et al.* 2017), making Iso-Seq an ideal technology for reconstructing a transcriptome without a reference genome sequence and without assembling fragments to resolve the complete isoform sequence (Honaas *et al.* 2016; Ju *et al.* 2016). Moreover, alternative transcripts that contribute to the gene space complexity (Brett *et al.* 2002) and vary with cell type (Swarup *et al.* 2016), developmental stage (Thatcher *et al.* 2016), and stress (Yan *et al.* 2012; Liu *et al.* 2016) cannot be definitively characterized without full-length transcript information.

The objective of this study was to test whether full-length cDNA sequencing with Iso-Seq technology is an effective strategy for reconstructing a grape transcriptome reference for transcriptional profiling without an annotated genome assembly. We compared how Cabernet Sauvignon’s Iso-Seq transcriptome fares as a reference for RNA-seq analysis versus its annotated genome. We sequenced the full-length transcripts of ripening berries with Iso-Seq and Illumina RNA-seq reads. The high-coverage short-read data were used to profile gene expression and to error-correct low-expression isoforms that would have been otherwise lost by the standard Iso-Seq pipeline. The transcriptome reference built with Iso-Seq data represented most of the expressed genes in the grape berries and included cultivar-specific or “private” genes. When used as the reference for RNA-seq, Iso-Seq generated transcriptome profiles quantitatively similar to those obtained by mapping on a complete genome reference. These results support using Iso-Seq to capture the gene space of a plant and build a comprehensive reference for transcriptional profiling without a pre-defined reference genome.

## METHODS

### Plant material and RNA isolation

Grape berries from Cabernet Sauvignon FPS clone 08 were collected in Summer 2016 from vines grown in the Foundation Plant Services (FPS) Classic Foundation Vineyard (Davis, CA, USA). Between 10 and 15 berries were sampled at pre-véraison, véraison, post-véraison, and at commercial maturity. **Table S1** provides weather information for the sampling days. The ripening stages were visually assessed based on color development and confirmed by measurements of soluble solids (**Figure S1; Table S2**). On the day of sampling, berries were deseeded, frozen in liquid nitrogen, and ground to powder (skin and pulp). Total RNA was isolated using a Cetyltrimethyl Ammonium Bromide (CTAB)-based extraction protocol as described in Blanco-Ulate *et al.* (2013). RNA purity was evaluated with a Nanodrop 2000 spectrophotometer (Thermo Scientific, Hanover Park, IL, USA). RNA was quantified with a Qubit 2.0 Fluorometer using the RNA broad range kit (Life Technologies, Carlsbad, CA, USA). RNA integrity was assessed using electrophoresis and an Agilent 2100 Bioanalyzer (Agilent Technologies, Santa Clara, CA, USA). Only RNA with integrity number (RIN) greater than 8.0 was used for SMRTbell library preparation.

### Library preparation and sequencing

RNAs from four biological replicates per developmental stage were pooled in equal amounts. One μg of the pooled RNA was used for cDNA synthesis and for SMRTbell library construction using the SMARTer PCR cDNA synthesis kit (Clontech Laboratories, Inc. Mountain View, CA, USA). First-strand cDNA synthesis was performed using the SMRTScribe Reverse Transcriptase (Clontech Laboratories, Inc. Mountain View, CA, USA). Each developmental stage was individually barcoded (**Table S3**). To minimize artifacts during large-scale amplification, a cycle optimization step was performed by collecting five 5 μl aliquots at 10, 12, 14, 16, and 18 PCR cycles. PCR reaction aliquots were loaded on an E-Gel pre-cast agarose gel 0.8 % (Invitrogen, Life Technologies, Carlsbad, CA, USA) to determine the optimal cycle number. Second-strand cDNA was synthesized and amplified using the Kapa HiFi PCR kit (Kapa Biosystems, Wilmington, MA, USA) with the 5’ PCR primer IIA (Clontech Laboratories, Inc. Mountain View, CA, USA) following the manufacturer’s instructions. Large-scale PCR was performed using the number of cycles determined during the optimization step (14 cycles). Barcoded double-stranded cDNAs were pooled at equal amounts and used for size selection. Size selection was carried out with a BluePippin (Sage Science, Beverly, MA, USA) and 1-2 kbp, 2-3 kbp, 3-6 kbp, and 5-10 kbp fractions were collected. After size selection, each fraction was PCR-enriched prior to SMRTbell template library preparation. cDNA SMRTbell libraries were prepared using 1-3 μg of PCR enriched size-selected samples, followed by DNA damage repair and SMRTbell ligation using the SMRTbell Template Prep Kit 1.0 (Pacific Biosciences, Menlo Park, CA, USA). A second size selection was performed on the 3-6 kbp and 5-10 kbp fractions to remove short contaminating SMRTbell templates. A total of 8 SMRT cells were sequenced on a PacBio Sequel system (DNA Technologies Core, University of California, Davis, USA). Demultiplexing, filtering, quality control, clustering and polishing of the Iso-Seq sequencing data were performed using SMRT Link ver. 4.0.0 (**Table S4**). Iso-Seq read error rates were estimated using the identity of the best alignment on the diploid Cabernet Sauvignon genomic assembly (Chin *et al.* 2016). Alignment was performed with GMAP ver. 2015-09-29 (Wu and Watanabe 2005) using the parameters “-K 20000 -B 4 -f 2”. Coding sequences (CDS) were identified using Transdecoder (Haas *et al.* 2013) as implemented in the PASA ver. 2.1.0 (Haas *et al.* 2003).

RNA-seq libraries were prepared using the Illumina TruSeq RNA sample preparation kit v2 (Illumina, San Diego, CA, USA), following the low-throughput protocol. Each biological replicate was barcoded individually. Libraries were evaluated for quantity and quality with the High Sensitivity chip on a Bioanalyzer 2100 (Agilent Technologies, Santa Clara, CA, USA) and sequenced on an Illumina HiSeq4000 (DNA Technologies Core Facility, University of California, Davis, USA; **Table S5**). Quality filtering and adapter trimming were performed with Trimmomatic ver. 0.36 (Bolger *et al.* 2014) using the following parameters: “ILLUMINACLIP:2:30:10 LEADING:7 TRAILING:7 SLIDINGWINDOW:10:20 MINLEN:36” (**Table S5**). Error correction of the Full-length Non-Chimeric (FLNC) Iso-Seq reads using was carried out using high-quality Illumina reads and LSC ver. 2.0 (Au *et al.* 2012) with a minimum coverage threshold of 5 reads (“--short_read_coverage_threshold 5”).

### Transcriptome reconstruction and annotation

Isoforms identified by the SMRT Link pipeline and error-corrected Iso-Seq reads were merged using EvidentialGene (Gilbert 2013) in order to obtain a non-redundant transcriptome (ISNT). Contaminant sequences were searched by parsing blastn (Altschul *et al.* 1990) alignments against the NCBI NT database (ftp://ftp.ncbi.nlm.nih.gov/blast/db/, retrieved January 17th, 2017) using Megan ver. 6.6.5 (Huson *et al.* 2007) with default parameters. Sequences detected as non-viridiplantae were removed. Isoforms with no RNA-seq read mapping on their sequence over the 16 samples (662 isoforms) were considered as putative artifacts and were also discarded. ISNT sequences are provided in **File S1**. Functional annotation was performed with blastx (Altschul *et al.* 1990) using RefSeq plant proteins as database (ftp://ftp.ncbi.nlm.nih.gov/refseq, retrieved January 17th, 2017) imposing an HSP length cutoff of 50 amino acids. Functional domains were identified with InterProScan ver. 5.28-68.0 (Jones *et al.* 2014) (**File S2)**. Tree view of identified GO terms was generated using WEGO ver. 2.0 (Ye *et al.* 2006). Iso-Seq reads were considered derived from repetitive regions when showing a RepeatMasker (Smit *et al.* 2013) hit with coverage ≥ 75% and an identity ≥ 50% using the custom created repeat library (see below). Non-coding RNAs were identified with Infernal ver. 1.1.2 (Nawrocki *et al.* 2009) using the Rfam database ver. 12.2 (Nawrocki *et al.* 2015). Secondary overlapping alignments and structures with an *e*-value ≥ 0.01 were rejected. Hits on the minus strand of the Iso-Seq reads were rejected as well as matches that were truncated or covering less than 80% of the entire read.

### Cabernet Sauvignon genome annotation

Cabernet Sauvignon primary contigs and haplotigs (Chin *et al.* 2016) were used as genomic reference. A repeat library was created *ad hoc* for Cabernet Sauvignon. MITEs were identified with MITEHunter ver. 11.2011 (Han and Wessler 2010); LTRs and TRIMs were identified with LTRharvest (GenomeTools ver. 1.5.7; Ellinghaus *et al.* 2008) and LTRdigest (GenomeTools ver. 1.5.7; Steinbiss *et al.* 2009). RepeatModeler ver. 1.0.8 (Smit and Hubley 2008), and RepeatMasker ver. open-4.0.6 (Smit *et al.* 2013) were used to generated a custom library of repeat models. Repetitive elements in Cabernet Sauvignon genome were then identified with RepeatMasker (Smit *et al.* 2013) using both custom and plant repeat models altogether (**Table S6**).

*Ab initio* trainings and predictions were carried out with SNAP ver. 2006-07-28 (Korf 2004), Augustus ver. 3.0.3 (Stanke *et al.* 2006), GeneMark-ES ver. 4.32 (Lomsadze *et al.* 2005), GlimmerHMM ver. 3.0.4 (Majoros *et al.* 2004), GeneID ver. 1.4.4 (Parra *et al.* 2000) and Twinscan ver. 4.1.2 (Korf *et al.* 2001; Brent 2008) using TAIR10 annotation for Arabidopsis as informant species. MAKER-P ver. 2.31.3 (Campbell *et al.* 2014a) was used to integrate the *ab initio* predictions with the experimental evidence listed in **Table S7** using the parameters reported in **File S3**. Only MAKER-P models showing an Annotation Edit Distance (AED) < 0.5 were kept.

Gene structure refinement was carried out with PASA ver. 2.1.0 (Haas *et al.* 2003), parameters can be found in **File S4.** As transcriptomic evidence, we used the Iso-Seq data as well all the publicly available grape transcriptomic data (**Table S7**). Public Cabernet Sauvignon RNA-seq data (**Table S7**) were *de novo* assembled separately *de novo* using HISAT2 and Stringtie ver. 1.1.3 (Pertea *et al.* 2015). *De novo* transcript sequences were then clustered in a non-redundant dataset using CD-HIT-EST ver. 4.6 (Li and Godzik 2006) with an identity threshold of 99%. Genome annotation summary is reported in **Table S8**. Alternative splicing forms were classified using AStalavista ver. 3.0 (Foissac and Sammeth 2007).

Non-coding RNAs were searched with Infernal ver. 1.1.2 (Nawrocki *et al.* 2009) as described above (**Table S9)**. The predicted transcripts’ functional annotation was made with blastp search using the RefSeq plant protein database. Functional domain identification was done with InterProScan as described above for the ISNT. For each gene locus, a non-redundant list of the GO terms attributed to all the alternative transcripts was generated.

### Gene expression analysis with RNA-seq

For expression profiling, short reads were aligned on reference sequences using Bowtie2 ver. 2.26 (Langmead and Salzberg 2012) with options “--sensitive --dpad 0 --gbar 99999999 --mp 1,1 --np 1 --score-min L,0,-0.1”. Evaluation of expression at isoform and gene locus levels was carried out using RSEM ver. 1.1.14b3 (Li and Dewey 2011) with default parameters. Differential expression analysis was performed using EBSeq ver. 1.16.0 (Leng *et al.* 2013). For each pairwise comparison of consecutive growth stages, size factors were calculated with median normalization using five iterations of the EM algorithm. Genes were considered as significantly differentially expressed if they had a minimum *posteriori* estimate probability (PPDE) threshold of 0.95 and mean RPKM greater than 1 in at least one of two growth stages. Pearson correlation matrices and PCAs were performed using the log_2_-transformed RPKM values. Heatmaps of the Pearson correlation matrices and dendrograms were generated using heatmap.2 function from the gplots R package ver. 3.0.1 (Warnes *et al.* 2016). PCAs were carried out in R with the FactoMineR package ver. 1.41 (Lê *et al.* 2008).

In order to compare expression values of ISNT transcripts and gene loci in the Cabernet Sauvignon genes, ISNT transcripts were aligned to the Cabernet Sauvignon genome using GMAP as described above. ISNT transcripts were associated with gene loci annotated on the Cabernet Sauvignon genome if they aligned at least for 66% of their length. Alignments with translocation were excluded (**Table S10**). For each ISNT-gene locus association, gene expression was calculated as the sum of the mean expression values of all ISNT transcripts and all Cabernet Sauvignon gene loci, separately, using RSEM ver. 1.1.14b3 (Li and Dewey 2011). Differential expression analysis at cluster level was performed using EBSeq ver. 1.16.0 (Leng *et al.* 2013).

## RESULTS

### Isoform sequencing of the grape transcriptome during berry development

To obtain a comprehensive representation of the transcripts expressed during berry development, we isolated RNA from Cabernet Sauvignon berries (**Figure 1**) before the onset of ripening (4.35 ± 0.39 °Brix), at véraison (10.94 ± 0.26 °Brix), after véraison (18.38 ± 0.61 °Brix), and at commercial ripeness (20.33 ± 0.76 °Brix). To avoid loading bias, cDNAs were fractionated based on their length to produce four libraries at each developmental stage in size ranges of 1-2 kbp, 2-3 kbp, 3-6 kbp, or 5-10 kbp (**Figure 1**). Libraries derived from different developmental stages were barcoded and libraries with similar cDNA size were pooled together. Each library pool was sequenced independently on two SMRT cells of a Pacific Biosciences Sequel system generating a total of 23.6 Gbp. In parallel, the same samples were sequenced using Illumina technology (25,655,771 ± 3,512,980 high-quality reads per sample) to provide high-coverage sequence information for error correction and for gene expression quantification (**Table S5**). Demultiplexing, filtering and quality control of SMRT sequencing data were performed using SMRT Link as described in the Methods section. A total of 672,635 full-length non-chimeric (FLNC; **Figure 2**) reads with a maximum length of 14.6 kbp and a N50 of 3.5 kbp were generated (**Table S4**). FLNC reads were further polished and clustered into 46,675 single representatives of expressed transcripts (henceforth, polished-clustered Iso-Seq reads or PCIRs) ranging from 400 bp to 8.8 kbp with a N50 of 3.6 kbp (**Table S4**). The alignment of FLNC reads and PCIRs to the genomic DNA contigs of the same Cabernet Sauvignon clone (Chin *et al.* 2016; Minio *et al.* 2017) confirmed that sequence clustering and polishing successfully increased sequence accuracy, whose median values were 95.4% in FLNC reads and 99.6% in the PCIRs. The increase in sequence accuracy was also reflected by the significantly longer detectable coding sequences (CDS) in the PCIRs compared to the short and fragmented CDS found in the FLNC reads (**Figure 2**). The residual sequence discrepancy between PCIRs and the genomic contigs could be explained by heterozygosity and/or sequencing errors, but unexpectedly not by expression level (**Figure S2**).

**Figure 1:**
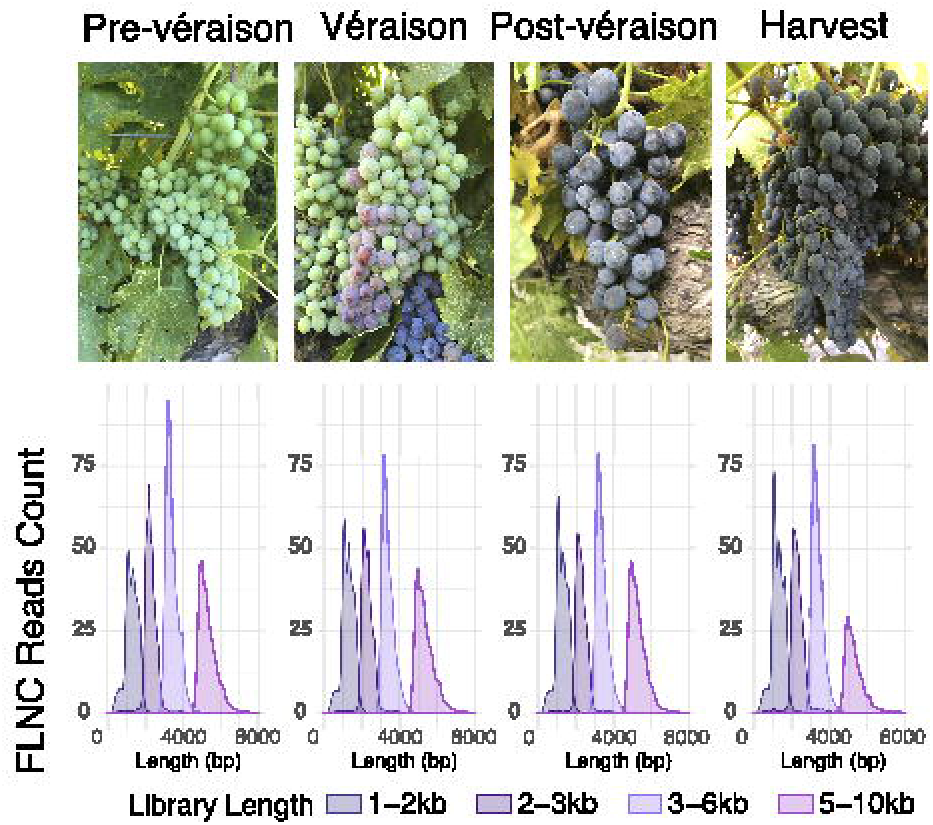
Full-length cDNA sequencing (Iso-Seq) of Cabernet Sauvignon berries. Representative pictures of berry clusters at the four growth stages used in this study and the read length distribution of the Iso-Seq libraries by cDNA size fraction.

**Figure 2:**
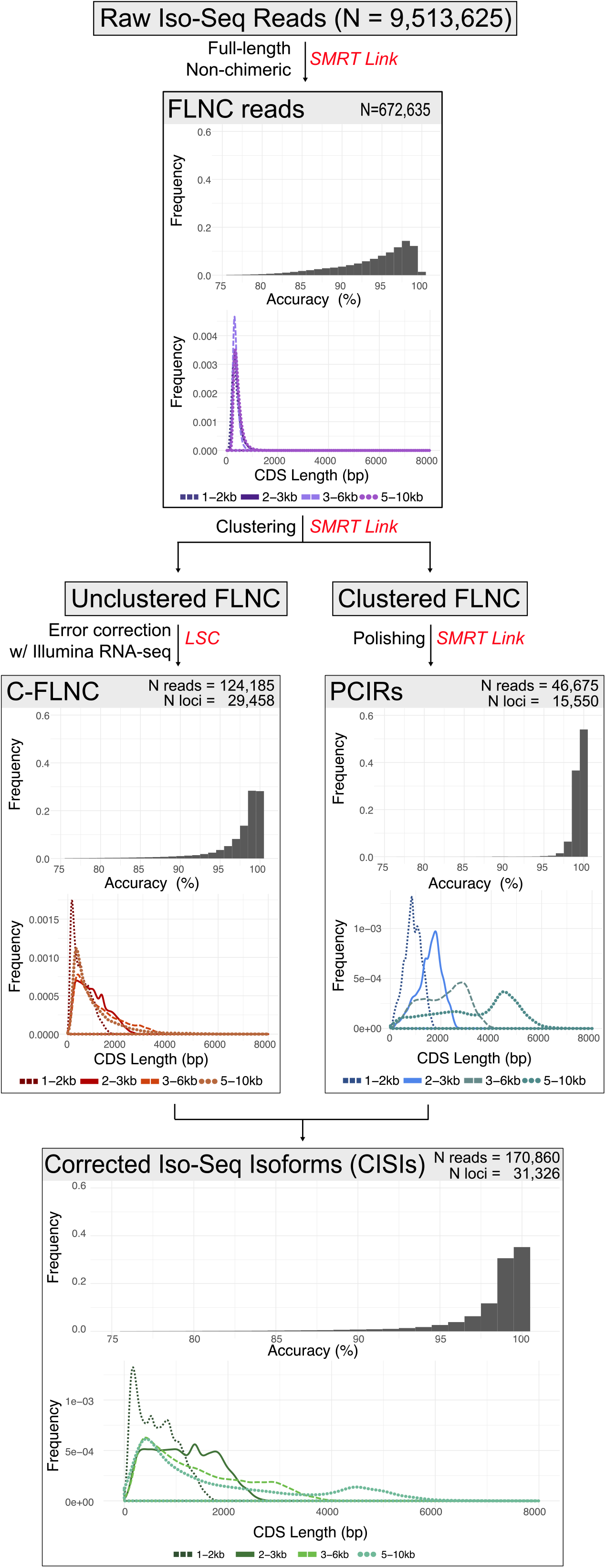
Diagram depicting the main steps of Iso-Seq read analysis. Raw Iso-Seq reads were processed to obtain Full-Length Non-Chimeric (FLNC) reads and clustered isoform reads (PCIRs). FLNC reads that did not cluster in any of the PCIRs were error-corrected using RNA-seq data (C-FLNC). The final dataset described in this study included PCIRs and C-FLNC reads. For each step, sequencing accuracy and CDS length distributions are reported.

Over 18.5% of the FLNC reads did not cluster with any other reads and were discarded by the SMRT Link pipeline. When mapped on the genomic contigs, the uncorrected reads displayed a sequence accuracy that reflected the typical error rate of 10 - 20% of the technology (**Figure 2**) (Giordano *et al.* 2017; Zimin *et al.* 2017; Koren *et al.* 2017). High error rates also resulted in short and fragmented detectable CDS (**Figure 2**). To recover the information carried by these 124,185 uncorrected FLNCs, which represented an important fraction of the transcriptome (see below), we error-corrected their sequences with LSC (Au *et al.* 2012) using the short reads generated with Illumina technology. As for the PCIRs, error correction resulted in greater sequence accuracy and longer CDS (**Figure 2**). PCIRs and error-corrected FLNC (C-FLNC) reads were finally combined into a single dataset of 170,860 corrected Iso-Seq isoforms (CISIs). As low as 1.7 % (2,826) of the CISIs showed significant homology with interspersed repeats. LTRs and LINEs were the most abundant orders with 778 and 729 representatives, respectively. Chloroplast and mitochondria genes represented a small fraction of the CISIs with only 89 isoforms (0.05%) having a significant match (50% identity and mutual alignment coverage). CISIs were also searched for non-coding RNA (ncRNA) using the covariance models of the Rfam database; only 182 isoforms were annotated as ncRNAs and were all ribosomal RNA (145 attributed to the large subunit, clan CL00112 and 37 to the small subunit, clan CL00111). Excluding these transcribed isoforms, only 164 CISIs (0.1%) failed to align to the Cabernet Sauvignon genomic contigs, confirming the completeness of the genome reference and the negligible biological contamination of the berry samples.

To reconstruct an Iso-Seq non-redundant transcriptome (ISNT) that would be tested as a reference for expression profiling, CISIs were clustered with EvidentialGene (Gilbert 2013) to reduce the redundancy between stages and libraries. One representative for each expressed transcript was retained for a total of 28,721 isoforms (**File S1**). Less than four percent of these isoforms did not contain a complete open reading frame, likely due to residual errors in their sequence. Mapping on the Cabernet Sauvignon genome showed that they potentially derive from 24,378 gene loci over both haplotypes (**File S5**). Most of isoforms (61.2%) aligned to two gene loci, one on each haplotype (i.e. primary and haplotig), as a result of the diploid phasing of the Cabernet Sauvignon genome assembly. For 10,727 loci (44%), multiple isoforms (2.03 ± 3.49 isoforms/locus) mapped on the same locus possibly due to alternative splicing and structural differences between alleles. With 77.6% of the BUSCO orthologous genes represented, the completeness of the ISNT was remarkably high, close to that of the entire Cabernet Sauvignon genome (96.7%; Chin *et al.* 2016), even if the ISNT was constructed using expression data from berries only. Interestingly, 301 BUSCO complete gene models (20%) were found in multiple copies in the ISNT, suggesting that alternative isoforms of these highly conserved genes are expressed during ripening. Putative functions were assigned to the ISNT transcripts as described in the Methods section (**Table S11**). Only four sequences did not match any sequence in the databases used. Gene Ontology (GO) terms were assigned to 23,386 transcripts with an average of ~6.9 GO terms per transcript (**Table S11, File S6-S8**).

### Isoform sequencing allows the discovery of private Cabernet Sauvignon genes

Previous analyses of gene content in a limited number of grape cultivars showed that up to 10% of grape isoforms were not shared between genotypes. Some of these “dispensable” genes were associated with cultivar-specific characteristics (Da Silva *et al.* 2013). To identify protein-coding transcripts characteristic of Cabernet Sauvignon (i.e., private genes), we looked for homologous sequences among the ISNT transcripts in the PN40024 genome (Jaillon *et al.* 2007; Vitulo *et al.* 2014) and in the transcriptomes of Corvina (Venturini *et al.* 2013), Tannat (Da Silva *et al.* 2013), and Nebbiolo (Gambino *et al.* 2017). Approximately five percent of the ISNT (1,501 isoforms) did not have a homologous copy in any of the three datasets (**Table S12**). These Cabernet Sauvignon private isoforms were involved in various biological processes of berry development and ripening like phenylpropanoid/flavonoid biosynthesis (a chalcone synthase, a flavanone 3-hydroxylase, and a flavonoid 3’-hydroxylase (Falginella *et al.* 2012)), sugar accumulation and transport (ten sucrose-phosphate synthases, a phosphofructokinase, a glucose-6-phosphate dehydrogenase, a sucrose transport SUC4-like, a polyol transporter, and an inositol transporter (Afoufa-Bastien *et al.* 2010; Xin *et al.* 2013)), water transport (eleven aquaporins), and cell wall metabolism and loosening (six cellulose synthases, a xyloglucan galactosyltransferase, one xyloglucan glycosyltransferase 9-like, two expansins, two xyloglucan endotransglucosylase/hydrolases, two pectinesterases, a pectin methylesterase/invertase inhibitor, seven polygalacturonases, and two ß-galactosidases (Carey *et al.* 1995; Cosgrove 2000, 2005)).

### Protein-coding gene models in the Cabernet Sauvignon genomic assembly

To evaluate the non-redundant Iso-Seq transcriptome’s completeness and usefulness as a reference for RNA-seq analysis, the protein-coding genes in the Cabernet Sauvignon genome were predicted as described in **Figure S3**. First, the repetitive regions of the genome were masked using a custom-made library of Cabernet Sauvignon MITE, LTR, and TRIM information. Overall, ~51% of the assembly consisted of repetitive elements (**Table S6**), with 412,994 repetitive elements on the primary assembly (313 Mb) and 274,123 on the haplotigs (177Mb), LTRs were the most abundant class, covering over 335 Mb of the genomic sequences, with Gypsy and Copia families accounting for 201 Mb and 104 Mb, respectively. Next, MAKER-P (Campbell *et al.* 2014b) identified putative protein-coding loci, combining the results of six *ab initio* predictors trained *ad hoc* with publicly available experimental evidences. *Ab initio* predictors were trained using a custom set of 4,000 randomly selected gene models out of the 5,636 high-quality, non-redundant, and highly conserved gene models of the PN40024 V1 transcriptome (4,459 multiexonic and 1,177 monoexonic). Experimental evidence from public databases (**Table S7**) was incorporated and used to validate the predicted models. The final MAKER-P prediction included 38,227 high-quality gene models (AED < 0.5) on the primary contigs and 26,789 on the associated haplotigs. Using the covariance models from the Rfam database, 5,780 non-overlapping putative long non-coding RNAs (ncRNAs; 3,239 on primary contigs and 2,541 on haplotigs) belonging to 275 different families were annotated (**Table S9**). Gene models were further improved using the information from all Iso-Seq full-length datasets (PCIRs, C-FLNC, FLNC), RNA-seq, and the publicly available grapevine transcriptome assemblies. This final refinement improved the annotation of the UTRs and added isoform information. PCIRs helped identify 155 new loci not detected by MAKER-P, update the structure of 10,801 gene models, and add 2,712 alternative transcripts. C-FLNC reads introduced 830 additional missing loci and added 3,738 alternative transcripts to the annotation. Together, 14,388 gene models were updated. FLNC reads introduced 14 new loci and 20,493 alternative transcripts, bringing the number of updated model structures to 24,945. Predicted genes without similarity to known proteins in the RefSeq database and without any functional domains identified by InterProScan (Jones *et al.* 2014) were removed. The final predicted transcriptome included 55,887 transcripts on 36,689 loci on primary contigs and 40,444 transcripts on 25,479 loci on haplotigs (**Table S8**). GO terms were assigned to 80,752 transcripts (83.8%) based on homology with protein domains in RefSeq and InterPro databases **(Table S13, File S9-S11**). A genome browser for Cabernet Sauvignon, its annotation, and an associated blast tool are available at http://cantulab.github.io/data.

For 2,995 (11%) of the 25,479 protein-coding gene loci identified on the haplotigs, we could not find a corresponding homolog in the primary contigs, likely reflecting the diversity in gene content between parental genomes (Cabernet Franc and Sauvignon Blanc (Bowers and Meredith 1997)), as in Corvina (2,321 transcripts; Venturini *et al.* 2013), Tannat (1,873 transcripts; Da Silva *et al.* 2013), and Nebbiolo (3,961 transcripts; Gambino *et al.* 2017). We could not find homologous genes in PN40024, Corvina, Tannat, and Nebbiolo for 1,714 protein-coding gene loci (**Table S14**). Those genes included likely members of the phenylpropanoid/flavonoid biosynthesis pathway: four phenylalanine ammonia-lyase, three chalcone synthases, a chalcone isomerase, a dihydroflavonol-4-reductase, a flavanone 3-hydroxylase, a flavonoid 3’,5’-hydroxylase, a leucoanthocyanidin dioxygenase, and an anthocyanin acyltransferase. Other private Cabernet Sauvignon genes were associated with terpenoid biosynthesis, including six valencene synthases that may play a role in grapevine flower aroma, three vinorine synthases (Lücker *et al.* 2004; Martin *et al.* 2009), and a (E,E)-geranyllinalool synthase. The incorporation of Iso-Seq data in the gene prediction pipeline also allowed the structural annotation of alternative transcripts. Twenty five percent (15,509) of the 62,168 annotated gene loci had two or more alternative isoforms, an average of 1.55 ± 1.29 alternative transcripts per locus, confirming previous reports in PN40024 (Vitulo *et al.* 2014). The frequency of splicing variant types was similarly observed in other plant species (Reddy *et al.* 2013). Intron retention was the most abundant type, accounting for over 44% (**File S12**), similar to what was observed for rice (45-55%) (Zhang *et al.* 2015), Arabidopsis (30 - 64%) (Marquez *et al.* 2012; Reddy *et al.* 2013; Zhang *et al.* 2015) and maize (40 - 58%) (Zhang *et al.* 2015; Wang *et al.* 2016). Alternative acceptor sites (13%), alternative donor sites (10%), and exon skipping (8%) were the other most abundant types of alternative splicing found in the Cabernet Sauvignon genome; a full description of the selected splicing events is reported in **File S12**.

### Iso-Seq transcriptome as reference for RNA-seq analysis

The final step of the analysis was to evaluate the effectiveness of the reconstructed ISNT as reference for RNA-seq analysis of berry development compared to the gene space predicted on the Cabernet Sauvignon genome. Comparisons between the two references are summarized in **Table 1**. Only about three percent more RNA-seq reads mapped on the Cabernet Sauvignon predicted transcriptome (90.6 ± 0.8 %) than on the ISNT (87.2 ± 0.8 %), suggesting that Iso-Seq reconstructed most of the transcripts detectable by RNA-seq at a coverage of ~26 M reads / sample. Approximately 75% of the ISNT (21,494 transcripts) and ~49% of the predicted gene space (30,501 gene loci) was detected as expressed (mean RPKM ≥ 1) in at least one stage (**Figure 3A, Table S15-S16**). In both datasets, the number of expressed genes was slightly higher at pre-véraison stage than at later developmental stages, consistent with previous observations of ripening Cabernet Sauvignon berries (Fasoli *et al.* 2018). For both datasets, Pearson’s correlation matrix and Principal Component Analysis (PCA) showed a clear distinction between pre-véraison stage and the three ripening stages, as well as a stronger correlation between post-véraison and full-ripe berry transcriptomes (**Figure 4**), confirming the well-known transcriptional reprogramming associated with the onset of ripening (Fasoli *et al.* 2012; Massonnet *et al.* 2017a) and suggesting that similar global transcriptomic dynamics of berry development can be obtained using either Iso-Seq or the whole genome as reference. We then applied a sequence clustering approach to define associations between ISNT isoforms and gene loci to directly compare the expression values of each gene in the two references. Based on reciprocal overlap of the alignment, we were able to associate 25,306 ISNT transcripts with 26,873 gene loci in the Cabernet Sauvignon genome (**Table S10**). Gene expression levels measured on the two references were well-correlated (*R* = 0.92; *P*-value < 2.2 e-16; **Figure 3B**; **Tables S17-18**). Differential gene expression analysis identified 14,477 ISNT transcripts and 18,600 Cabernet Sauvignon genes significantly differentially expressed (PPDE ≥ 0.95) at least once during berry development (**Table S15-S16**). More genes were differentially regulated between pre-véraison and véraison than during ripening for both references, (**Figure 5A & B**) as previously observed (Palumbo *et al.* 2014; Massonnet *et al.* 2017a). Ninety one percent of the differentially expressed ISNT isoforms were also differentially expressed when RNA-seq data were mapped on genomic loci. Similar relative amounts of Biological Process GO terms among differentially expressed genes were observed between the two references (**Figure 5C & D**). Interestingly, 302 Cabernet Sauvignon private isoforms (transcripts not found in other cultivars) were differentially expressed during berry development, including transcripts encoding a polyol transporter, an inositol transporter, and five aquaporins.

**Figure 3:**
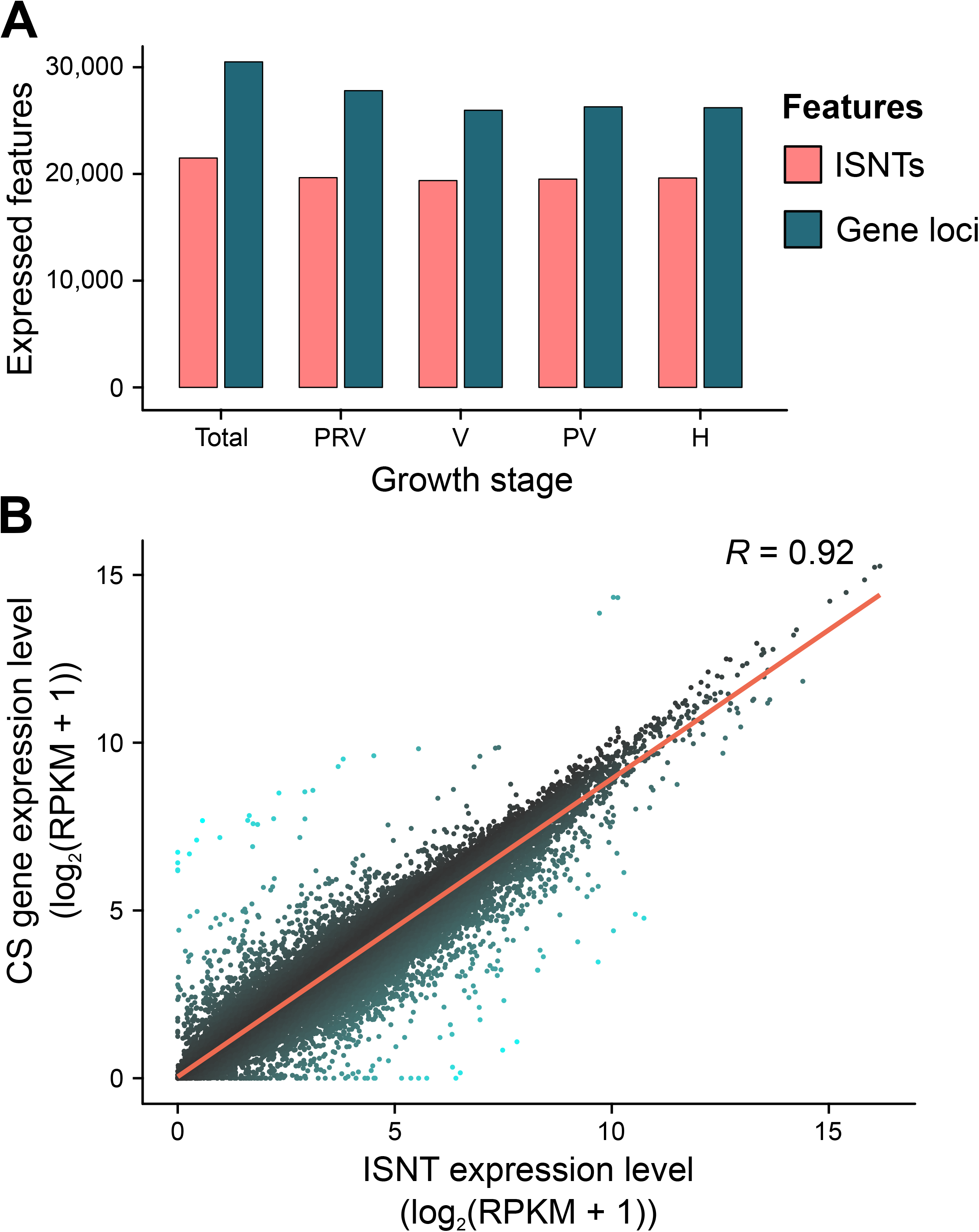
Gene expression analysis during berry development using the ISNT as reference and comparison with the Cabernet Sauvignon genome. (**A**) Number of ISNT transcripts (ISNTs) and gene loci expressed overall and at each ripening stage. (**B**) Scatterplot showing the correlation between the gene expression level (log_2_(RPKM + 1)) of ISNT isoforms and Cabernet Sauvignon (CS) gene loci. Line of best fit and correlation coefficient factor (*R*) are provided.

**Figure 4:**
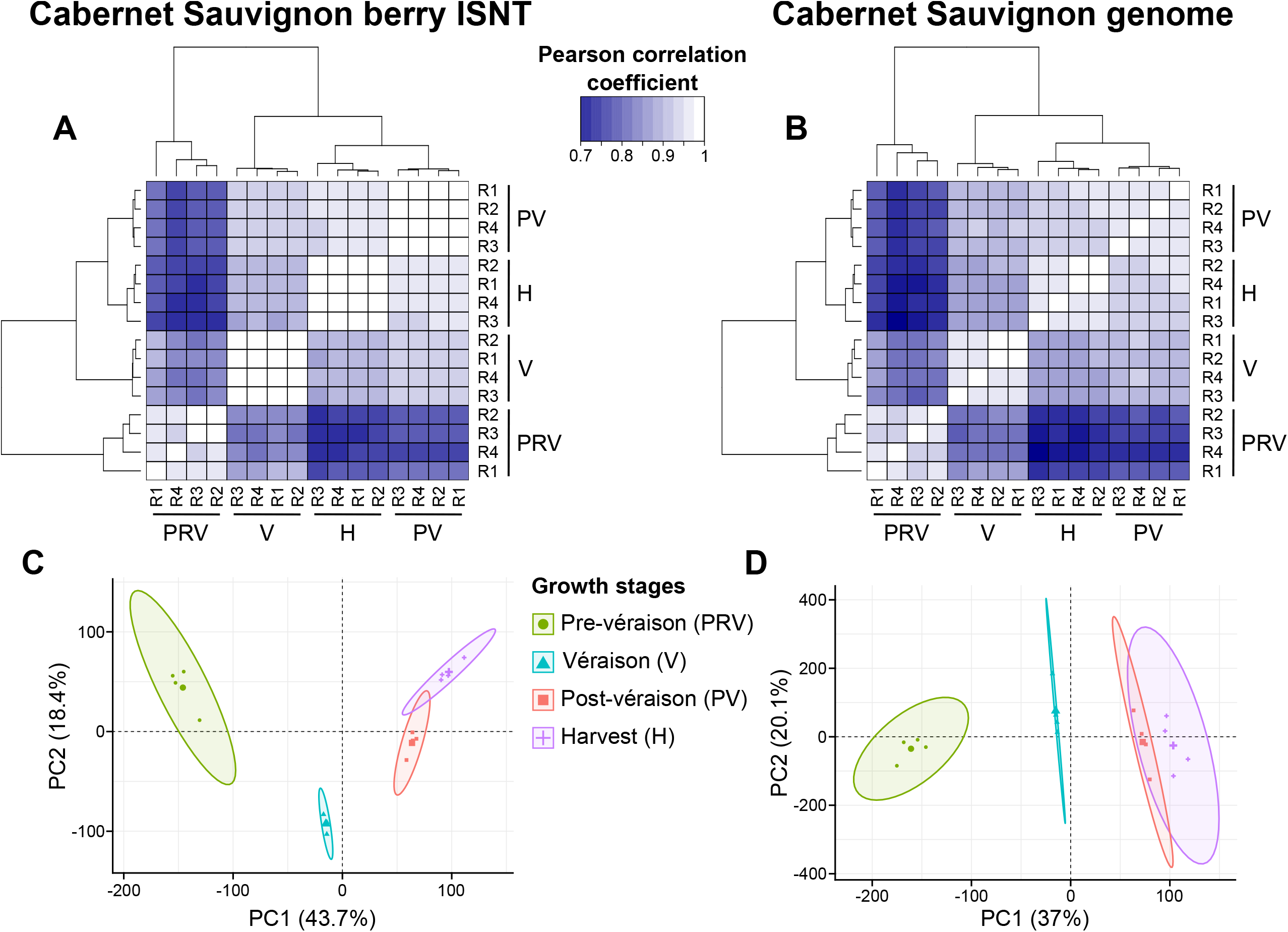
Global transcriptomic changes during berry development ripening using the ISNT and Cabernet Sauvignon gene space as references. (**A, B**) Pearson correlation matrices of the 16 berry transcriptomes. (**C, D**) PCA plots of the 16 berry transcriptomes. Each point represents a biological replicate and ellipses define confidence areas (95%) for each berry growth stage. All analyses were performed using the log_2_-transformed RPKM values of expressed features (mean RPKM ≥ 1 at least at one stage).

**Figure 5:**
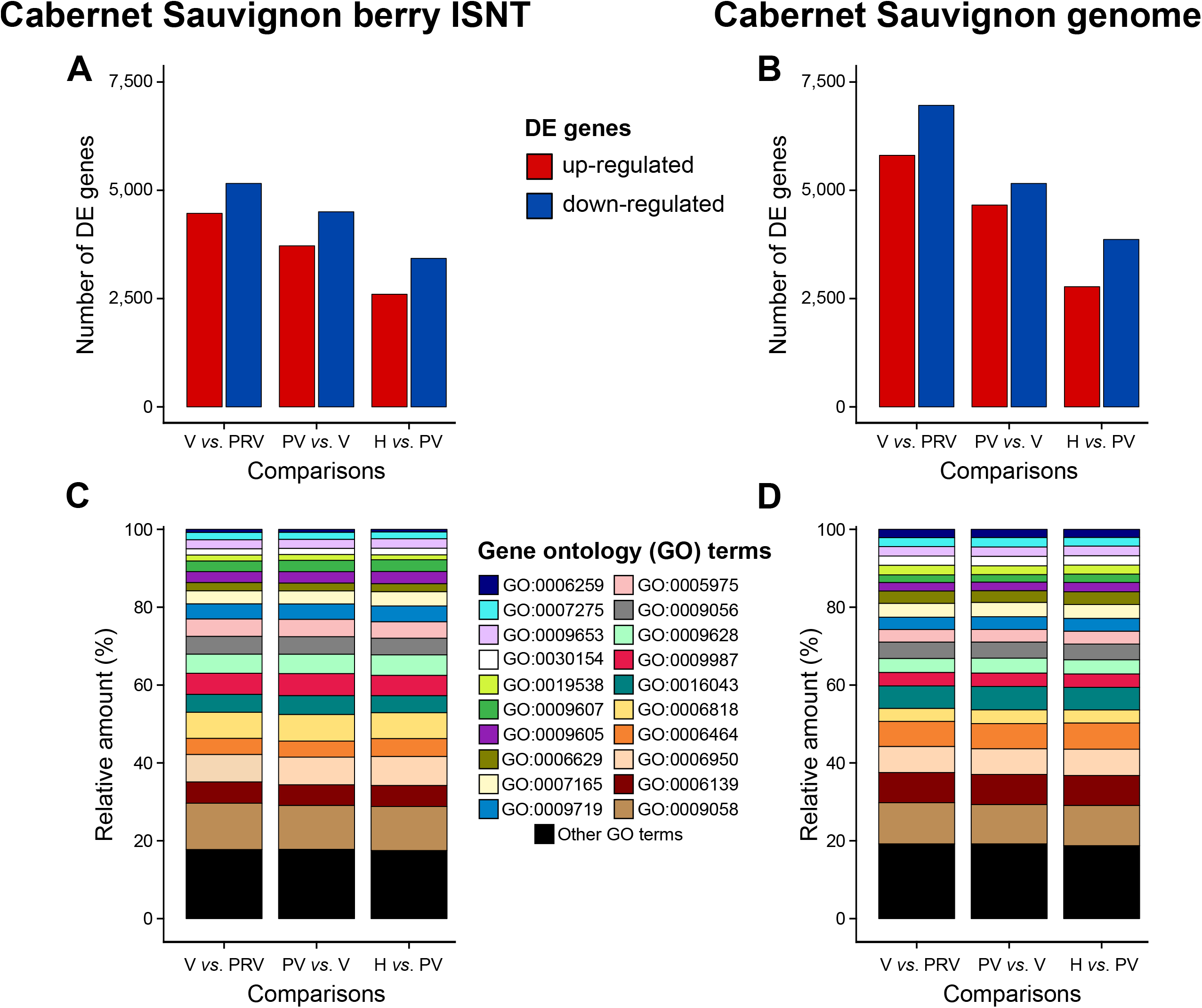
Comparison of the differential expressed ISNT transcripts and Cabernet Sauvignon gene loci during berry development. (**A, B**) Number of up- and down-regulated genes between each pairwise comparison of developmental stages (PPDE ≥ 0.95 and mean RPKM ≥ 1 at least at one stage). (**C, D**) Relative amount of Biological Process Gene Ontology (GO) terms among differentially expressed genes.

**Table 1:** Summary of the RNA-seq results when Cabernet Sauvignon berry ISNT and the Cabernet Sauvignon genome were used as reference for short-read mapping. *Percentage of ISNT; **percentage of Cabernet Sauvignon predicted transcriptome; Biological process, BP; differentially expressed, DE.

|  | Iso-Seq transcriptome | Cabernet Sauvignon genome |
| --- | --- | --- |
| Mapping rate | 87.2 ± 0.8 % | 90.6 ± 0.8 % |
| Expressed features (mean RPKM ≥ 1 at least at one growth stage) | 21,494 isoforms (74.8%)* | 30,501 gene loci (49%)** |
| Expressed features with BP GO term annotation | 14,431 isoforms | 21,588 gene loci |
| Total BP GO terms among expressed features | 61,603 GO terms | 80,103 GO terms |
| DE features (PPDE ≥ 0.95 and mean RPKM ≥ 1 at least at one growth stage) | 14,477 isoforms (50.4%)* | 18,600 gene loci (29.9%)** |
| DE features with BP GO term annotation | 10,237 isoforms | 14,026 gene loci |
| Total BP GO terms among DE features | 44,179 GO terms | 53,349 GO terms |

## DISCUSSION

Full-length cDNA sequencing with SMRT technology (Iso-Seq) can be used to rapidly reconstruct the grape berry transcriptome, enabling the identification of cultivar-specific isoforms, refinement of the Cabernet Sauvignon genome annotation, and the creation of a reference for transcriptome-wide expression profiling. In contrast to transcriptome reconstruction using short-read sequencing that requires *de novo* assembly, Iso-Seq delivers full-length transcripts that eliminate the introduction of assembly errors and artefacts like chimeric transcripts and incomplete fragments due to PolyA capture (Chang *et al.* 2014; Huang *et al.* 2016; Moreton *et al.* 2016; Smith-Unna *et al.* 2016; Geniza and Jaiswal 2017; Ungaro *et al.* 2017). The incorporation of high-coverage short-read sequencing is still necessary to benefit from the complete transcript sequencing enable by Iso-Seq. Although Iso-Seq provides much longer reads than second-generation sequencing platforms and as a result is excellent in resolving transcript structure, its sequencing error rate is high (10 - 20%) and throughput is still relatively low (Koren *et al.* 2016; Giordano *et al.* 2017; Zimin *et al.* 2017). Here we show that combining Iso-Seq with Illumina sequencing at high coverage enables expression profiling and sequence error correction of Iso-Seq reads, particularly those derived from low-expression genes. The clustering analysis of the SMRT link pipeline discarded ~18.5% of the FLNC reads, likely caused by low sequence accuracy. To overcome this technical issue, we applied a hybrid error correction pipeline consisting in performing the error-correction of the unclustered FLNC reads, followed by an additional clustering step of both to resolve redundancies. Error correction with Illumina reads recovered a significant amount of Iso-Seq reads that would have otherwise been removed by the standard Iso-Seq pipeline, highlighting the importance of integrating multiple sequencing technologies with complementary features (Koren *et al.* 2012; Au *et al.* 2012; Salmela and Rivals 2014; Hu *et al.* 2016).

Transcriptome reconstruction has been widely used to develop references for genome-wide expression profiling in the absence of an annotated reference genome assembly (Simon *et al.* 2009; Garber *et al.* 2011; Martin and Wang 2011; Yang and Kim 2015). Though a genome reference is available for grape, transcriptome reconstruction overcomes the limitations of a cultivar-specific reference that lacks the gene content of other cultivars. Although cultivar-specific genes appear nonessential for berry development, those private genes could contribute to cultivar characteristics. For example, the wine grape Tannat accumulates unusual high levels of polyphenols in the berry; cultivar-specific genes account for more than 80% of the expression of phenolic and polyphenolic compound biosynthetic enzymes (Da Silva *et al.* 2013). *De novo* transcriptome assembly from short RNA-seq reads has been used to explore the gene content diversity in Tannat (Da Silva *et al.* 2013), Corvina (Venturini *et al.* 2013), and Nebbiolo (Gambino *et al.* 2017). Iso-Seq identified 1,501 Cabernet Sauvignon transcripts expressed during berry development that were found in neither the genome of PN40024 nor the transcriptomes of Tannat, Nebbiolo and Corvina. Some private Cabernet Sauvignon transcripts have functions potentially associated with traits characteristic of Cabernet Sauvignon grapes and wines like their color and sugar content. These transcripts included three sugar transporter-coding genes, which could be involved in the accumulation of glucose and fructose during berry ripening (Lecourieux *et al.* 2014), and a chalcone synthase, a flavanone 3-hydroxylase, and a flavonoid 3’-hydroxylase, all involved in the flavonoid pathway. Chalcone synthases catalyze the first committed step of the flavonoid biosynthesis pathway (Sparvoli *et al.* 1994), which produces different classes of metabolites in grape berry, including flavonols (yellow pigments), flavan-3-ols and proanthocyanidins (mouth-feel and smooth sensory perceptions), and anthocyanins (red, purple, and blue pigments). In addition, products of the flavonoid 3’-hydroxylase can lead to the synthesis of cyanidin-3-glucoside, a red anthocyanin (Castellarin *et al.* 2012). The analysis of the gene space in the genome assembly showed that private Cabernet Sauvignon genes identified using Iso-Seq are only a fraction of the private Cabernet Sauvignon transcriptome. As in other transcriptome reconstruction methods, Iso-Seq can only identify transcripts that are expressed in the organs and developmental stages used for RNA sequencing. Obtaining the full set of private transcripts without genome assembly would require sequencing additional organs and developmental stages. In addition, it is challenging to differentiate isoforms derived from close paralogous genes, alleles of the same gene, and alternative splicing variants, in any transcriptome obtained by RNA sequencing (including Iso-Seq); this potentially leads to an overestimation of the genes in the final transcriptome reference. This study could not resolve isoform redundancy in the final transcriptome for about 37% of the gene loci in the Cabernet Sauvignon genome. This is a limitation of Iso-Seq as well as of all transcriptome references that cannot be overcome without a complete genome assembly.

In this study, we tested whether the transcriptome reconstructed using Iso-Seq can be used for expression profiling. Only an approximately 3% difference in read alignment between ISNT and the genome reference was observed, implying that at high-coverage, ISNT detects almost all genes expressed during berry development. The slight difference in mapping rate between the two references can be explained by either the absence of some low-expression transcripts in the ISNT or the residual error rate in isoform sequences. Gene expression analysis using the ISNT as reference showed similar results compared to the Cabernet Sauvignon genome assembly, with a very high correlation of expression level and differential gene expression, and with similar global transcriptomic changes. However, we observed differences in the number of expressed and differentially expressed features that depend on the reference used. Those differences could be explained by the diploid phasing of the Cabernet Sauvignon genome assembly and that multiple ISNT transcripts might correspond to a single gene locus. Nonetheless, similar relative amounts of Biological Process GO terms were found among the differentially expressed genes, confirming that the transcriptome obtained using Iso-Seq captured the transcriptional reprogramming underlying the main physiological and biochemical changes during grape berry development. In addition, gene expression analysis revealed that some private isoforms (20%) are significantly modulated during berry development, indicating that in addition to identifying the private gene space, the ISNT reference makes it possible to observe its expression.

In conclusion, this study demonstrates that Iso-Seq data can be used to create and refine a comprehensive reference transcriptome that represents most genes expressed in a tissue undergoing extensive transcriptional reprogramming during development. In grapes, this approach can aid developing transcriptome references and is particularly valuable given diverse cultivars with private transcripts and accessions that are genetically distant from available genome references, like the non*-vinifera Vitis* species used as rootstocks or for breeding. This pipeline described here can be useful in efforts to reconstruct the gene space in plant species with large and complex genomes still unresolved.

## Acknowledgments

This work was supported by J. Lohr Vineyards and Wines, E. & J. Gallo Winery, and the Louis P. Martini Endowment in Viticulture.

## List of abbreviations

FLNC: full-length non-chimeric
PCIR: polished-clustered Iso-Seq read
C-FLNC: error-corrected FLNC
CISI: corrected Iso-Seq isoform
ISNT: Iso-Seq non-redundant transcriptome

## Data Availability

Sequencing data are accessible through NCBI (**PRJNA433195**) and other relevant datasets, such as protein-coding gene and repeat coordinates, can be retrieved from the Cantu lab github repository (http://cantulab.github.io/data.html).

File S1: Iso-Seq reconstructed transcriptome (FASTA format)

File S2: databases used for InterProScan search of functional domains

File S3: parameters used for MAKER-P annotation

File S4: parameters used in PASA for annotation polishing

File S5: association of Iso-Seq reconstructed transcripts with gene loci in the Cabernet Sauvignon genome

File S6: biological process GO tree of the Iso-Seq reconstructed transcriptome

File S7: cellular component GO tree of the Iso-Seq reconstructed transcriptome File S8: molecular function GO tree of the Iso-Seq reconstructed transcriptome

File S9: biological process GO tree of the protein-coding genes predicted in the Cabernet Sauvignon genome

File S10: cellular component GO tree of the protein-coding genes predicted in the Cabernet Sauvignon genome

File S11: molecular function GO tree of the protein-coding genes predicted on the Cabernet Sauvignon genome

File S12: distribution and classification of alternative splicing events annotated on the Cabernet Sauvignon genome

Figure S1: Soluble solids content of Cabernet Sauvignon berries at four different growth stages. For each biological replicate, soluble solid measurement (°Brix) was performed using two technical replicates.

Figure S2: Evaluation of the impact of the expression level on Iso-Seq sequence accuracy. Density plot describing the distribution of PCIRs (polished-clustered Iso-Seq reads) based on the measured sequence accuracy and the cumulative expression value measured using RNA-seq.

Figure S3: Cabernet Sauvignon genome annotation pipeline. The diagram represents the workflow used to produce the gene annotation of Cabernet Sauvignon genome.

Table S1: Weather conditions during the sampling of Cabernet Sauvignon berries.

Table S2: Soluble solids content (°Brix) of Cabernet Sauvignon berries at the four developmental stages.

Table S3: Sequences of the oligo dT barcodes used for the construction of the Iso-Seq SMRTBell libraries.

Table S4: Iso-Seq read sequencing and standard PacBio clustering pipeline statistics.

Table S5: Short-read sequencing, filtering and mapping results.

Table S6: Repetitive content identification statistics.

Table S7: Experimental evidences used for MAKER annotation.

Table S8: Gene and transcript annotation statistics.

Table S9: RFAM categories identified in Cabernet Sauvignon genome.

Table S10: Groups of ISNT isoforms and annotated gene loci based on sequence clustering approach.

Table S11: Functional annotation of the reconstructed ISNT.

Table S12: Functional annotation of the cultivar-specific ISNT isoforms.

Table S13: Functional annotation of the Cabernet Sauvignon genome.

Table S14: Functional annotation of the Cabernet Sauvignon private gene loci.

Table S15: RNA-seq analysis of the 16 berry samples using the Iso-Seq non-redundant transcriptome (ISNT) as reference. Evaluation of expression was carried out using RSEM ver. 1.1.14b3 (Li and Dewey 2011). Raw expression values are provided, as well normalized expression values and RPKM values. Differential expression analysis was performed using EBSeq ver. 1.16.0 (Leng et al. 2013) for each pairwise comparison of consecutive growth stages. Abbreviation of the four developmental stages: Pre-véraison, PRV; Véraison, V; Post-véraison, PV; Harvest, H.

Table S16: RNA-seq analysis of the 16 berry samples using the Cabernet Sauvignon genome as reference. Evaluation of expression was carried out using RSEM ver. 1.1.14b3 (Li and Dewey 2011). Raw expression values are provided, as well normalized expression values and RPKM values. Differential expression analysis was performed using EBSeq ver. 1.16.0 (Leng et al. 2013) for each pairwise comparison of consecutive growth stages. Abbreviation of the four developmental stages: Pre-véraison, PRV; Véraison, V; Post-véraison, PV; Harvest, H.

Table S17: Expression and differential expression analysis of the clustered ISNT transcripts. Evaluation of expression was carried out using RSEM ver. 1.1.14b3 (Li and Dewey 2011). Raw expression values are provided, as well normalized expression values and RPKM values. Differential expression analysis was performed using EBSeq ver. 1.16.0 (Leng et al. 2013) for each pairwise comparison of consecutive growth stages. Abbreviation of the four developmental stages: Pre-véraison, PRV; Véraison, V; Post-véraison, PV; Harvest, H.

Table S18: Expression and differential expression analysis of the clustered Cabernet Sauvignon gene loci. Evaluation of expression was carried out using RSEM ver. 1.1.14b3 (Li and Dewey 2011). Raw expression values are provided, as well normalized expression values and RPKM values. Differential expression analysis was performed using EBSeq ver. 1.16.0 (Leng et al. 2013) for each pairwise comparison of consecutive growth stages. Abbreviation of the four developmental stages: Pre-véraison, PRV; Véraison, V; Post-véraison, PV; Harvest, H.

